# *Plasmodium* circumsporozoites targeted by RFdiffusion / cyclic-peptides

**DOI:** 10.1101/2025.09.01.673429

**Authors:** J Coll

## Abstract

The surface-exposed circumsporozoite-protein (CSP) of blood-migrating *Plasmodium falciparum*, is the target for preventive vaccines inducing 70-80 % protection against human malaria. Rather than for prevention, therapeutic-like peptides computationally designed to recognize full-length CSP were generated here aiming to complement the present vaccines. To generate thousands of lineal-peptide backbones and amino acid sequences, most recent deep-learning RFdiffusion algorithms were employed to target CSP including its disordered repeats. Optimal affinities were best predicted by several conformers generated from 34-mer lineal-peptides. Additional cyclic-peptide sequences, affinities and more rigid conformer structures were predicted by Alphafold2-cyclization. Thus easy-to-synthesize representative-top cyclic-peptides predicting picoMolar affinities were generated targeting CSP domains including hepatocyte-binding, NANP disordered-repeats and/or cell-adhesion. The cyclic-peptide results of applying these novel computational strategies to full-length CSP could be employed for additional experimental validation.

## *Plasmodium* CSP

The protozoan *Plasmodium falciparum* cause human malaria by mosquito’s inoculation of 10-100 elongated-gliding unicellular circumsporozoites per bite^1^. Blood circumsporozoites migrate into the liver to replicate to release thousands of rounded merozoites per hepatocyte cell^2^. Back into the blood, merozoites infect erythrocytes ^3^ for additional replication/amplification/ to generate clinically-detectable, deathly infections^4^.

Most of the external membranes of **<u>C</u>**ircum**<u>s</u>**porozoites are densely coated by one **<u>p</u>**rotein (CSP) of 397 amino acids. CSP accounts for 5–15 % of all circumsporozoite proteins. Each CSP molecule contains an N-terminal α-helix hydrophobic **<u>s</u>**ignal **<u>p</u>**eptide (SP) for membrane insertion into the circumsporozoite surface^2^, which is shortly removed in mature CSP. Each mature surface CSP molecule contains an N-terminal **h**eparan **s**ulfate **<u>p</u>**roteo**<u>g</u>**lycan (HSPG) sequence to bind hepatocytes ^5, 6, 7^ followed by a protease cleavage site for CSP activation and a large central domain of protective 3D-disordered repeats (129-273 residues^16^). Five amino acids (asparagine, alanine, valine, aspartate and proline) form disordered NANP, NPDP and NVDP repeats. A C-terminal part codes for a GPI membrane-anchor (**<u>g</u>**lycosyl-**<u>p</u>**hosphatidyl**i**nositol), a cell-adhesive TSR (**<u>t</u>**hrombo**<u>s</u>**pondin **r**epeat) and a terminal hydrophobic α-helix transmembrane (TM). Compared to ∼ 200 protein homologous of different species, *P*.*falciparum* TSR has unique sequences and disulphide bonds^8^.

During circumsporozoite migration in human blood, the CSPs maintain a non-adhesive cell conformation masking their TSR domains to avoid recognition by host antibodies. Upon finding hepatocyte cell membranes, both N- and C-terminal CSP domains contribute to recognition and binding^5, 6,7^. Thus, N-terminal HSPG hepatocyte-binding triggers the CSP proteolytic cleavage, unmasks the C-terminal TSR, and release CSP adhesive conformations to begin invasion^7^. Interferences with these circumsporozoite CSP conformational changes have been the target for vaccine prevention and for possible therapeutic drugs.

Previous intensive work demonstrated that antibodies against CSP N-terminal, NANP repeats and/or TSR were protective in mice models^9^ and were induced also by most recent human vaccines^10, 11^. Comparison of ∼ 200 monoclonal antibodies (mAb) developed by mice-immunization with purified circumsporozoites, showed that only those mAbs that included nanoMolar affinities toward NANP repeats could inhibit malaria infections^12-16^. In particular, the mAb850 induced NANP ß-turn conformations in the otherwise disordered NANP 3x and 6x repeats. Complexes of 19 mAb850 Fabs induced larger spirals in disordered NANP 28x repeats (conditional-folding?) correlating with their picoMolar affinities (i.e., models: 7uym.pdb, 7v05.pdb) ^17, 18^. Furthermore, mAb850 inhibited *P*.*falciparum* both *in vitro* and infected mice^18, 19^. In contrast, to anti-NANPs, antibodies recognizing TSR were rare, most probably due to their masking before CSP proteolytic cleavage^20^.

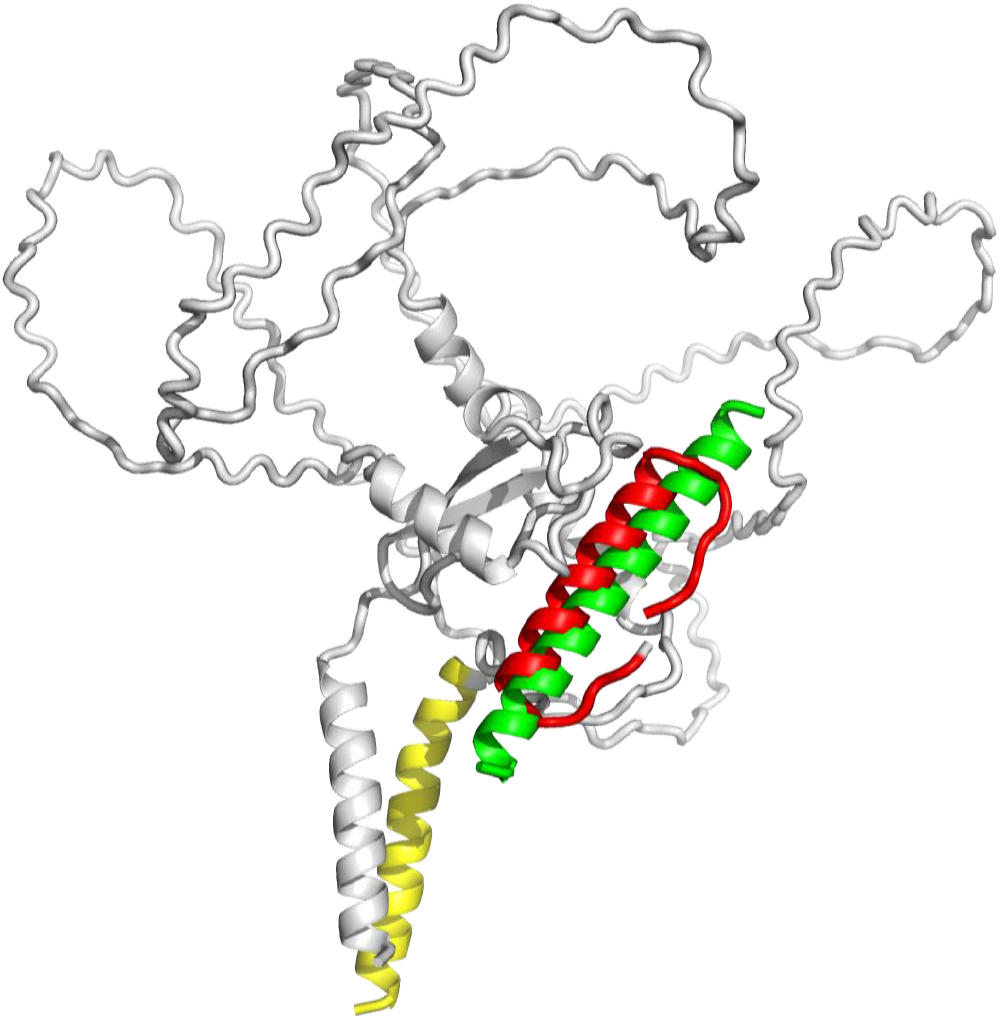

After numerous investigations, a vaccine inducing ∼ 40 % protection, was first developed by including CSP 19 NANP repeats and TSR (RTS,S) ^33^.

During recent years anti-*Plasmodium* vaccines have been further fine-tuned and approved to combat malaria (RTS,S and R21)^10, 11^ by the World Health Organization (WHO)^19,37^. These included the most recent R21/Matrix-M vaccine predicting 70-80 % protection^20-22^ against children clinical malaria across several African countries^22, 23^.

Complementing vaccines^31, 32^ and/or some reported therapeutic anti-CSP antibodies^24^, small drug-like molecules, either by traditional screening or by computational docking have also targeted circumsporozoite CSP. For instance, because only the CSP C-terminal domain 3D-structure could be solved by crystallization^27^, some previous computational docking efforts were recently reported targeting those domains ^25^. Rather than screening large molecular banks^26, 27^, computationally-mimicked co-evolution^25^ was employed to generate thousands of drug-like compounds. Compared to the screening of large banks, the co-evolution strategy generated deeper and faster penetration into the vast drug-like chemical space^6, 7^, as reported in other protein-ligand pairs^28-37^. However, although targeting C-terminal CSP^25^, hundreds of non-toxic^8^, nanoMolar affinity drug-like candidates, were generated, their often difficult-to-synthesize molecular structures challenged their experimental validation possibilities.

In contrast, small peptides facilitate their chemical or biological synthesis and therefore may offer easier access to experimental validation. Because of targeting external-membrane, CSP cyclic-peptides have the additional advantage of avoiding possible membrane permeability difficulties (which may pose additional problems in other cases)^38-45^. For these reasons, cyclic-peptides may add new tools for anti-malaria validation research. Therefore, peptides were explored by targeting full-length CSP of *P. falciparum* circumsporozoites.

After trial-and-error and feed-backs, a computational strategy consisting in two generation/selection cycles was developed. Step-by-step, the designed strategy consists in:

1. **Lineal-peptide generations**. RFdiffusion random generation of thousands of lineal-peptides, backbones and sequences, targeting a full-length CSP Alphafold2 model (following previous experiences ^25, 37, 46, 47^),
2. **Lineal-peptide selection**. Focus on the best size predicting top affinities,
3. **Cyclic-peptide generations**. Random generations of five cyclic-peptide models per lineal-peptide by Alphafold2 batch-cyclizations, and
4. **Cyclic-peptide selections**. Selection of the top cyclic-peptide sequences predicting picoMolar affinities for synthesis and experimental validations.

## Computational results

### *De novo* lineal-peptides generated by RFdiffusion

To explore any surfaces by RFdiffusion, a CSP full-length model (chain A) was selected to be used as target without defining any hot-spots. RFdiffusion were tested by ML (**<u>M</u>**a**<u>L</u>**aria) 53 runs that generated 16 lineal-peptide backbones per run and n0-n5 sequences per backbone (chain B). Lineal-peptides were named by a unique label made by joining their ML number, their number of design and their n sequence (i.e., ML34.3n0, ML46.6n3, ML53.15n3).

Lineal-peptide sizes from 4 to 60-mer (4 to 60 amino acid lengths) were explored among other criteria (not shown). The 34-mer lineal-peptide size predicted the maximal nanoMolar affinities in the 10^4^-10^5^ pM range (not shown). Therefore, the studies focused on further RFdiffusion *de novo* generations of lineal-peptide backbones and sequences targeting full-length CSP by 34-mer peptide designs. Because of the random nature of RFdiffusion, ML runs were replicated (ML34, ML46 and ML53). The results confirmed the maximal nanoMolar affinities of the generated lineal-peptides backbones/sequences in the 10^4^-10^5^ pM range (**Figure 1, open circles, green, red, blue**).

**Figure 1.**
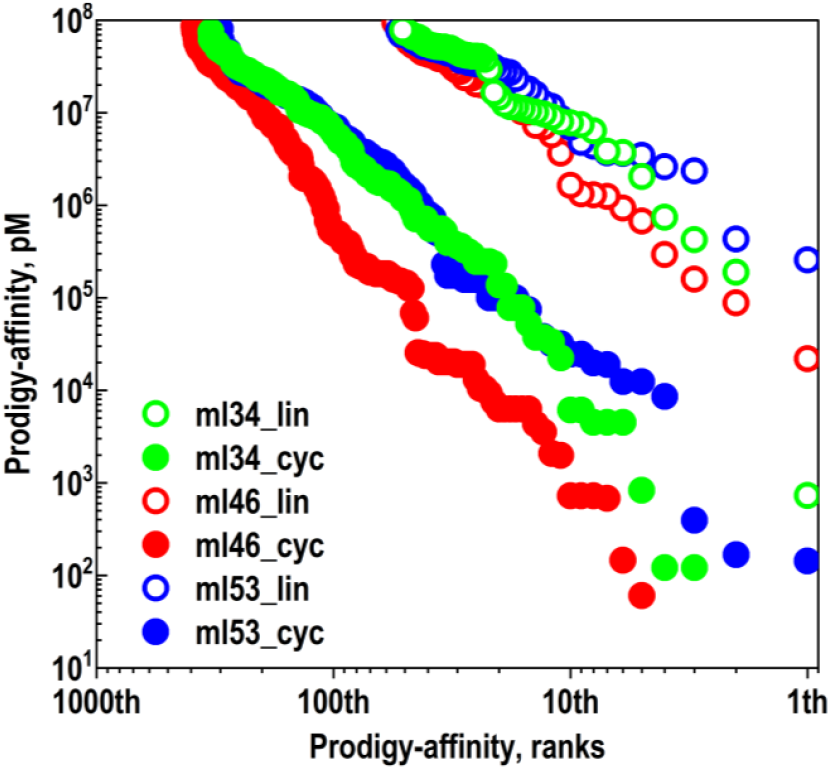
Affinity rank profiles of 34-mer peptides. *Plasmodium falciparum* 3D7-strain full-length CSP (PF3D7_0304600, XM_001351086) 1-397 residues Alphafold2 model was targeted. Three RFdiffusion+Cyclization runs (ML 34, ML46 and ML53) were performed from the optimal 34-mer lineal peptides targeting the full-length of CSP under the same conditions. Each RFdiffusion ML run generated 16 peptide backbone designs (0-15) with 6 amino acid sequences per design (n0-n5) (**Supplementary Materials / diffusionMain34-6.ipynb**). Cyclizations used batch-modified af_ cyc_design. ipynb ^48^,to generate _0 to _5 cyclic peptide sequences per lineal-peptide (**Supplementary Materials / af_cyc_design-JCM10.10batch.ipynb**). Affinity of the *de novo* generated peptide models were approximated by Prodigy. **Open-green circles**, ML 34 lineal-peptides. **Solid-green circles**, ML34 cyclic-peptides. **Open-red circles**, ML46 lineal-peptides. **Solid-red circles**, ML46 cyclic-peptides. **Open-blue circles**, ML53 lineal-peptides. **Solid-blue circles**, ML53 cyclic-peptides.

### Cyclic-peptides generated from lineal-peptides

Six cyclic-peptides were generated for each lineal-peptide backbone/sequence, to explore for possible affinity improvements by AF2-batch-cyclizations. The newly generated cyclic-peptides were labeled by adding a number to their name from _0 to _5. Prodigy affinities, pdb, amino acid sequences and contacts, were all predicted for each new chain A CPS + chain B cyclic-peptide complexes (A+B). The affinity rank profiles of the newly generated A+B complexes predicted higher affinities (lower scores) for most of the cyclic-peptide sequences, compared to their lineal-peptide counterparts (**Figure 1, closed** vs **open circles, green, red, blue**). Similar improvement of affinities by cyclization were confirmed in more than 30-other runs with different criteria (not shown). Despite the limited and relative accuracy of most predictions, these results also confirmed previous reports comparing cyclic-*versus* lineal-peptide affinities,^51^.

### Comparison between lineal- and top cyclic-peptides

Top cyclic-peptides were defined by filtering all the cyclic-peptides generated by RFdiffusion and AF2cyclization, by their unique amino acid sequences and < 9000 pM affinities (**Table1**). Confirming the rank profile data, the highest affinities (lower scores) were predicted by the top cyclic-peptide sequences, compared to their corresponding lineal-peptide counterparts. Many of the affinity prediction differences between cyclic- and their lineal-peptides, ranged > 1000-fold. These affinity improvements might be not only due to the newly acquired cyclic backbones but also to an better fitting of their new amino acid sequence contacts. On the other hand, the higher affinities of cyclic-peptides could be also explained by their more rigid conformers contributing to minimize docking energies, facilitating the random selection of amino acid new contacts.

**Table 1.**
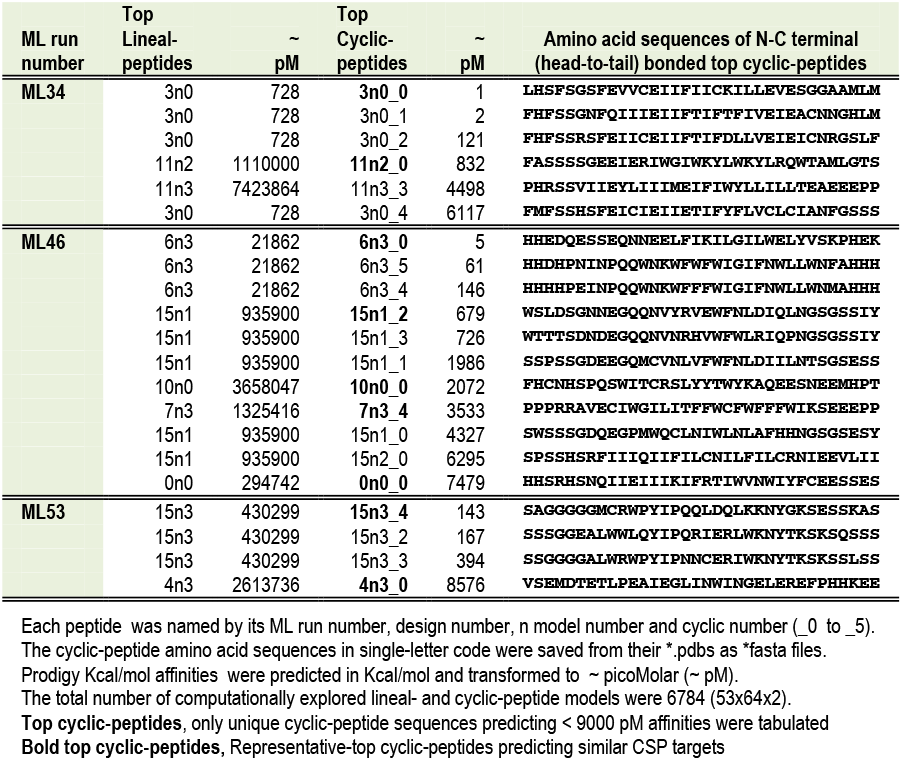
Affinity comparisons of top cyclic-peptides with their lineal-peptide counterparts.

### CPS amino acid contacts to top cyclic-peptides

CSP amino acid contacts to the top cyclic-peptides in the A+B complexes, predicted more similarity among those cyclic-peptides (labeled with _) that were generated from the same lineal-peptide (i.e., 3n0 lineal > 3n0_ cyclic-peptides, 6n3 lineal > 6n3_cyclic-peptides) (**Figure 2, same colors**). For instance:

**Figure 2.**
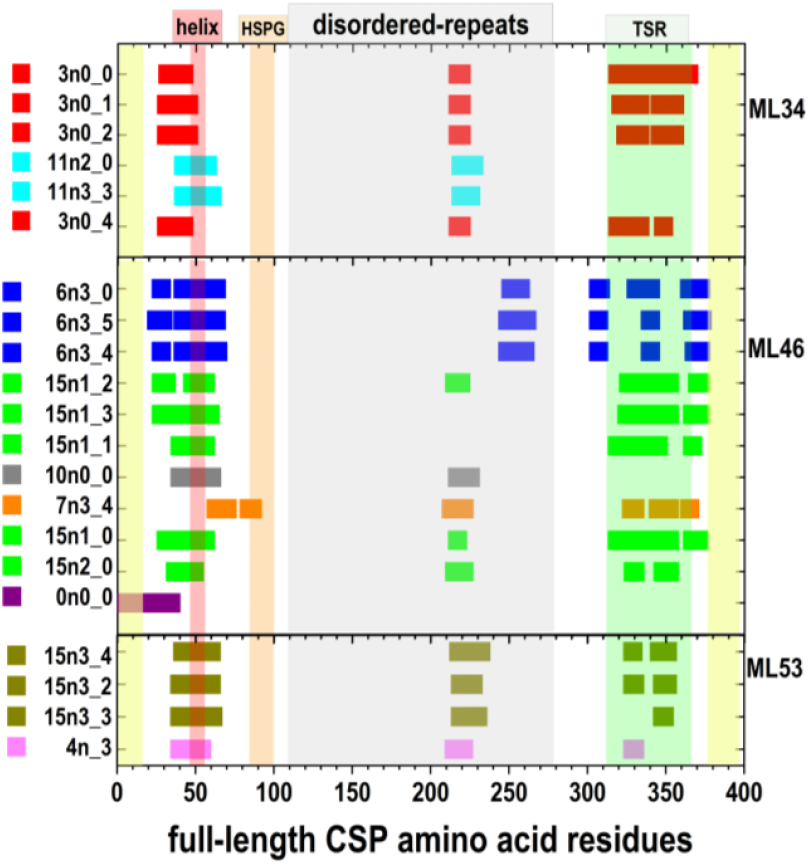
CSP amino acids nearby the top cyclic-peptides. The cyclic-peptides were generated as indicated above (**Figure 1** and **Table 1**). The CSP amino acids nearby the top cyclic-peptides were identified by Prodigy and graphically drawn in Origin. **Yellow SP**, Alphafold2-predicted signal peptide (1-26 residues). **Yellow TM**, Alphafold2-predicted transmembrane α-helix (376-395). **N-terminal helix**, Alphafold2-predicted α-helix (45-55)^49^. **HSPG**, hepatocyte surface binding (85-92) + Protease cleavage, ^93^KLKQP (93-97)^6^. **Disordered repeats:** 3xNANPNVDP, Alphafold2-predicted 3D disordered repeats (105-128) + 35x NANPs, Alphafold2-predicted 3D disordered repeats (129-272). **TSR short helix**, ∼TSR α-helix from 3vdj.pdb (312-324)^8.^ + cell adhesive, ∼ TSR domain from 3vdj.pdb (331-347)^8^ + protective segment, ∼ CS-flap from 3vdj.pdb (348-363)^24^ + disulphide bonds conserved among *Plasmodium* species (^338^C-^369^C and ^342^C-^374^C).

ML34.3n0 > ML34.3n0_ contacted the pre-amino-helix, ∼ 220 disordered repeats and TSR (**Figure 2, red**).

ML34.11n2 > ML34.11n2_ contacted the amino-helix and ∼ 220 disordered repeats (**Figure 2, cyan**).

ML46.6n3 > ML46.6n3_ contacted the CSP pre- and amino-helix, the ∼250-260 disordered repeats and around and inside TSR domain (**Figure 2, blue**). ML46.15n1 > ML46.15n1_ contacted the pre- and amino-helix, the ∼ 220 disordered repeats by some of the family and different parts mostly inside the TSR (**Figure 2, green**).

The ML46.10n0 > ML46.10n0_0, ML46.7n3 > ML46.7n3_4, and ML46.0n0 > ML46.0n0_0, each contacted some of the mentioned above CSP domains (**Figure 2, gray, orange, violet**, respectively). The ML46.7n3_4 was unique in contacting the post-amino helix and part of the HSPG region.

The ML53n3 > ML53.15n3_4 and the ML53.4n3 > ML53.4n3_0 contacted the amino-helix, the ∼210-230 disordered repeats and small parts of TSR (**Figure 2, dark green** and **light violet**, respectively).

The absence of contacts to SP (1-26 CSP residues), would suggest a reduced target for most top cyclic-peptides, except for ML46.0n0_0. On the other hand, top cyclic-peptides predicting picoMolar affinities, may target CSP during or shortly after CSP translation, before its SP removal and transfer to the circumsporozoite membrane, raising the question of whether or not they could even target circumsporozoite generation on mosquitos.

Some of the top cyclic-peptide included targeting of NANP repeats at the 220-230, 210-230 and/or 250-260 CSP residues (**Figure 2)**. Because NANP repeats have been immunodominant in malaria mouse models, human infections and/or protective vaccines, their recognition may suggest possible implications in protection by top cyclic-peptides. It might be encouraging that induction of *in vitro* neutralization, inhibition of hepatocyte invasion, and reduction of parasite numbers in mouse liver infections, have previously shown correlations with picoMolar binding to NANP repeats and similar affinities including NANP targetings have been predicted by some of the present top cyclic-peptides…….

### Visualization of CSP complexes with representative-top cyclic-peptides

To visualize A+B complexes by PyMol (**Figure 3** and **4**), representative-top cyclic-peptides were selected among those top cyclic-peptides predicting similar CSP targets (**Table 1, Bold top cyclic-peptides**). Most CSP domains were targeted by at least one of these representative-top cyclic-peptides:

**Figure 3.**
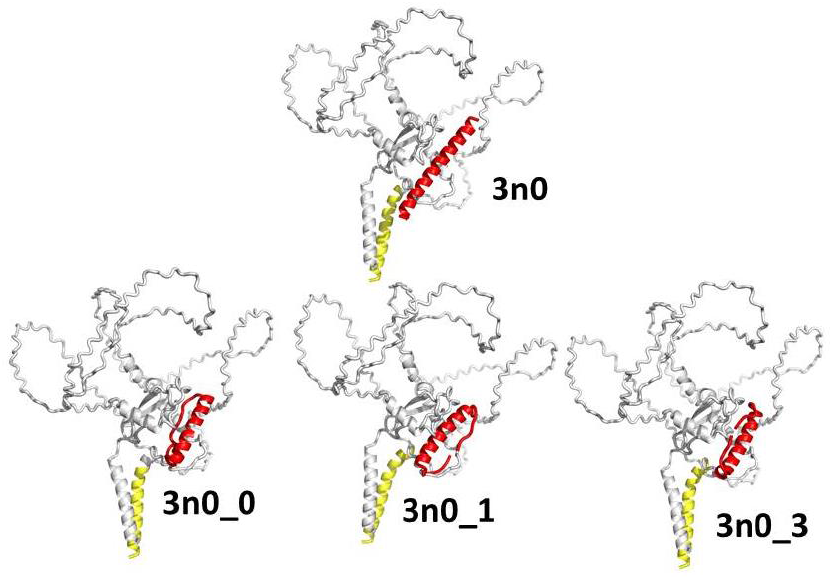
Comparison of 3n0 lineal and derived cyclic-peptides in complex with full-length CSP. The 3n0 lineal-peptide generated 6 cyclic-peptides (_0 to _5) by a home-batch modification of the Colab notebook af_ cyc_design.ipynb ^48^. Only the unique sequence cyclic-peptides were represented. **Yellow cartoons**, signal peptide SP α-helix (1-26 residues) of CSP. **Red cartoons**, 3n0 lineal- and top cyclic-peptides (drawn in PyMol as open cartoons but drawn closed when drawn as sticks (not shown).

**Figure 4.**
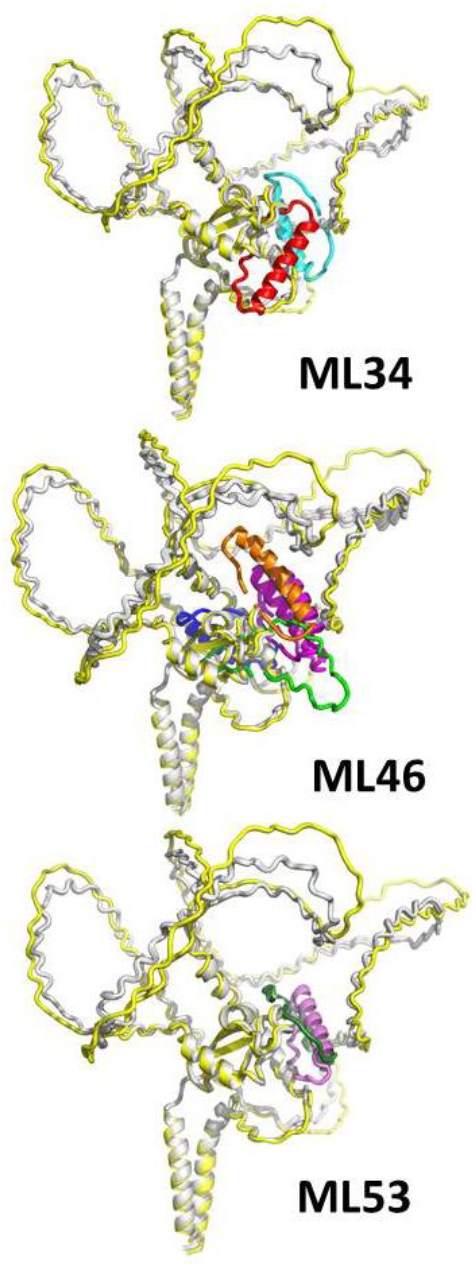
Representative-top cyclic-peptides complexed with full-length CSP. The RFdiffusion 34-mer lineal-peptides were derived to cyclic-peptides^48^ (**Table 1** and **Figure 2**). The most representative complexes of CSP with top cyclic-peptides of unique sequences < 200 pM affinities were represented here by PyMol. More details to explore at: **Supplementary Materials / ML34.pse**, **ML46.pse, ML53.pse**. **Yellow cartoons**, full-length CPS.pdb model from Alphafold2. **Gray cartoons**, merged CPS from the resulting complexes with representative-top cyclic-peptides (**Figure 2**). **ML34** **Red cartoon**, ML34.3n0_0 **Cyan cartoon**, ML34.11n2_0 **ML46** **Blue cartoon**, ML46.6n3_0 **Green cartoon**, ML46.15n1_2 **Gray cartoon**, ML46.10n0_0 **Orange cartoon**, ML46.7n3_4 **Violet cartoon**, ML46.0n0_0 **ML53** **Dark green cartoon**, ML53.15n3_4 **Light violet cartoon**, ML53.4n3_0

- ML34.3n0_0, ML34.11n2_0,
- ML46.6n3_0, ML46. 15n1_2, ML46.10n0_0, ML46. 7n3_4, ML46.0n0_0,
- ML53.15n3_4 and ML53.4n3_0.

An example comparing the lineal-peptide ML34.3n0 with some of its top cyclic-peptides (ML34.3n0_), illustrates their conformational changes (**Figure 3, up red cartoons**), despite predicting different backbones (lineal *vs* cyclic), amino acid sequences and affinities (**Table 1**). After cyclization, the ∼ N and C terminal 6 amino acids of the 3n0 lineal-peptide α-helix, were bended to each other as visualized for 3n0_0, 3n0_1, and 3n0_3 cyclic-peptides by acquiring extended structures (**Figure 3, down red cartoons**).

The visual comparison of the CSP complexes with representative-top cyclic-peptides, confirmed the coverage of the most important CSP domains, including the amino α-helix implicated in hepatocyte-binding (**Figure 4, orange vertical rectangle**), some of the NANP disordered-repeats (**Figure 4, gray vertical rectangle)** and/or the C-terminal TSR structure (**Figure 4, green vertical rectangle**).

Small conformational differences were detected between the full-length Alphafold2 CSP model (**Figure 4, yellow cartoons**) and the CSP complexed with representative-top cyclic-peptides, specially those mapped to disordered NANP repeats (**Figure 4, gray cartoons**). These conformational changes in the predicted 3D models suggest that the fixbb cyclization protocol not only changed the cyclic-peptide backbones and sequences (chain B), but also affected the CSP backbone conformations (chain A). The results suggest that both A+B conformations were optimized in a peptide-dependent manner. During fixbb cyclization, the input peptide backbone (chain B) is variable while the input CSP (chain A) mainly serves as a fixed target. However, during optimization the chain A also can undergo small conformational relaxations to optimize the interactions with the docked cyclic-peptides, specially those implicating the disordered NANP repeats. Nevertheless, most of their conformations in the complex remained close to the input structure due to their rigidity. The higher flexibility of the NANP disordered repeats suggest they would be more susceptible to conformational changes to improve cyclic-peptide docking.

### Computational limitations

Some of the main reasons to explore peptide cyclization were to reduce possible physiological degradation and to increase chemical stability. Because of their N-C terminal amide-links, head-to-tail cyclic-peptides are more resistant to physiological exopeptidase degradation, compared to their lineal-peptide counterparts. Therefore, cyclic-peptides should increase their persistence and activity in the host blood to best interfere with circumsporozoites. Additionally, the more rigid structure of cyclic-peptides compared to the lineal-peptides, reduces their chemical unfolding and/or aggregation, facilitating their transport and delivery at different temperatures. Although intracellular penetration is a known challenge for therapeutic cyclic-peptides > 30-mer^43-50^, in our present case their targeting external CSP would be permeability-independent.

The cyclic-peptides proposed here are limited to computational predictions. Therefore, practical conclusions may be influenced by simplification constrains applied (i.e., absence of water interactions, approximated affinites, CSP model dependence, possible resistant mutations, targeting a short number of CSP domains and/or other). However, despite those limitations, and before some of those other limitations are addressed, the top cyclic-peptide candidates described offer for the first time some computational possibilities to interfere with CSP functions. Their possible inhibition of *Plasmodium* virulence strictly depends on additional *in vitro* or *in vivo* experimental validations.

Because the random algorithms presently available are still undergoing fine-tuning, they required a large number of runs to identify any top peptide candidates. It is hoped that additional code, improving affinity predictions, including affinity filters during the generations, developing “genetic” alternatives, etc, could help to improve future explorations into the enormous peptide space^36, 39^

## Conclusions

The computational cyclic-peptides described predicted high affinities to *P*.*falciparum* external full-length CSP. Their CSP targets included both crystal-solved and some disordered-repeat domains, in contrast to previous drug-like compounds only targeting crystal-solved domains. Furthermore, compared to small drugs, peptides and top cyclic-peptides are easier-to-synthesize alternatives which predicted picoMolar rather than nanoMolar docking affinities. Because targeting externally coated *Plasmodium* circumsporozoites, cyclic-peptides should not require membrane permeability like other therapeutic cyclic peptides. These and other factors may work in combination, making peptide-cyclizations powerful strategies to improve potency and stability in peptide drug developments against *Plasmodium* infections.

## Computational methods

### *Plasmodium falciparum* circumsporozoite protein models

The 397 amino acid full-length **<u>c</u>**ircum**<u>s</u>**porozoite **<u>p</u>**rotein (CSP) sequence XM_001351086 of the *P*.*falciparum* reference 3D7 was alphafold modeled as described before ^45,25^.The most abundant similar representative of 10 alphafold models (CSP.pdb) was selected for docking. Only the C-terminal 301-396 residues, included their crystallized TSR-motif (3vdj.pdb)^7^, one GPI-anchor motif and the alphafold modeled transmembrane α-helix (**Figure 2**).

### Generation of lineal-peptides by RFdiffusion

Lineal-peptide were designed by **<u>R</u>**osetta**<u>F</u>**old diffusion (RFdiffusion)^50,51, 52^ (https://colab.research.google.com/github/sokrypton/ColabDesign/blob/main/rf/examples/diffusion.ipynb) accessed before the 15th of August of 2025. The RFdiffusion random noising/denoising cycles generated different amino acid backbones. Next ProteinMPNN generated best fitting amino acid for each backbone position taking into account user-added constrains. The final prediction structures were finally filtered by Alphafold2 ^51^. All RFdiffusion runs performed here maintained the same: contigs: A1-397, pdb: CPS.pdb, hotspot: ““, num_designs:16, add_potential: OK, use_beta_model: OK, num_seqs: 4, mpnn_sampling_temp: 0.3, run_aa: C, use_solubleMPNN: OK, initial_guess: OK, and num_recycles: 6 (**Supplementary Materials / diffusionMain34-6.zip**). Additional input variations included: the ranges of the CSP chain A contig residues to be targeted (i.e., A1-397:, A101-276: etc), and peptide-binder targeted size (amino acid number) (i.e., :34, :20:12, :40, etc). At the first cell, the RFdiffusion code was modified to include Driver mount and the final cell included automatically downloading-saving to Drive the zipped results. Once the results.zip were transferred into the local computer environment, two PyMol/Python scripts (RFana13.py and Prodigybatch21.py) were consecutively run to unzip several directories and order the lineal-peptides according to their Prodigy affinity estimations, respectively.

### Estimation of CSP-peptide complex affinities by Prodigy

Additionally to the Prodigy web service (https://wenmr.science.uu.nl/prodigy/), a local anaconda3-guided Prodigy-Python 3.10.8 installation at windows 10, was locally run including short and long interactions as recommended (“-- distance cutoff” of 5.5 Å^42^).The Windows-Prodigy version was run in batch as described before^34,46^ and modified here by PyMol/Python home-designed for-loop scripts (**Supplementary Materials / Prodigybatch21.py.zip)**. Prodigy algorithms^56-60^ estimated CSP-peptide complex affinities by validation of input pdb structures, calculation of interatomic interface distances, and buried surface area, by Biopython https://www.biopython.org), freesasa (https://github.com/mittinatten/freesasa), and naccess (http://www.ncbi.nlm.nih.gov/pubmed/994183), respectively. The Prodigy outputs included the following directories: all_pdb/ (for each of the pdb file complexes), all_contacts/ (a text file containing all predicted amino acid contacts), all_pml (PyMol pml scripts to draw all side chains in the contact surfaces), and all_Kcal (estimated average of total docking affinities in Kcal/mol). The home-script script also generates a file named all_kcalpM.txt (calculated picoMolar affinities by the formula 10^12*(exp(Kcal/0.592)), and an orderpdb/ directory (containing all_pdb ordered by the affinities calculated from all_kcalpM.txt).

### Cyclization of peptides

Head-to-tail cyclization of the linear RFdiffusion peptides (N-C terminal bonds) were performed in batch using a modified Colab notebook developed from the af_ cyc_design.ipynb ^48^ https://github.com/sokrypton/ColabDesign/blob/main/af/examples/af_cyc_design.ipynb. Code details were provided at **Supplementary Materials / af_cyc_design_JCM10.10batch**. A fixbb fixed backbone design protocol was employed and results automatically downloaded-saved at Drive. All the cyclic-peptide *.pdbs were submitted to Prodigy and PyMol as mentioned above.

## Supplementary information

**Table S1.**
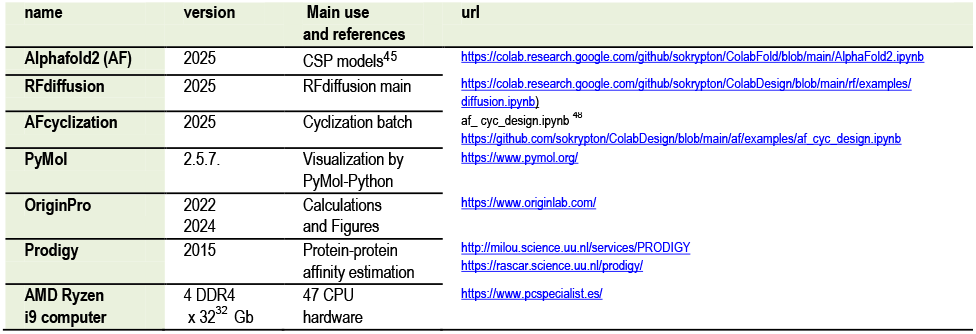
Computational software.

**Table S2.** Colab notebook-ipynb and Analysis-scripts.

| Generation designs | inputs | Generation *.ipynb | intermediate zip outputs | Analysis-scripts *.py | Outputs |
| --- | --- | --- | --- | --- | --- |
| I) RF-DIFFUSION | Contigs+ peptide size | DiffusionMain34-6. | *.results.zip | RFana13.<br>Prodigybatch21. | all_pdb/<br>ordered_pdb/<br>all_contacts/<br>all_pml/<br>all_KcalpM.txt |
| II) AF-CYCLIZATION | ordered_pdb | af_cyc_design_JCM10.10batch. | *batchpdb.zip | Prodigybatch21. | batchpdb/<br>all_pdb/<br>ordered_pdb/<br>all_contacts/<br>all_pml/<br>all_KcalpM.txt |

RF-DIFFUSION: https://colab.research.google.com/github/sokrypton/ColabDesign/blob/main/af_examples/diffusion.ipynb.
Slightly modified to Supplementary Materials / DiffusionMain34-6.
AF-CYCLIZATION: https://github.com/sokrypton/ColabDesign/blob/main/af_examples/af_cyc_design.ipynb<sup>48</sup>
Developed to Supplementary Materials / af\_cyc\_design\_JCM10.10batch
Both \*.ipynb notebooks start by Driver upload and end by automatic download/save their zipped results to Driver.
The Prodigy runs from a local Windows-installed Prodigy version adjusted to contacts within 5.5 Å.
Prodigybatch21.py runs as a subprocess of RFana13.py home scripts to facilitate analysis and affinity estimations.
\*.ML number of run.
The \*.py scripts are designed to run PyMol / Python by copy them into the same directory as their targeted files

## Supporting information

af_cyc_designJCM10_10batch.ipynb

diffusionMain34_6.ipynb

GraphycalAbstract

ML34.pse+ML46.pse+ML53.pse

Prodigybatch21

## Supplementary Materials

### GraphycalAbstract.zip

The full-length mRNA of *Plasmodium falciparum* reference CSP was modeled in Alphafold2. **Gray cartoons**, the most representative Alphafold2 predicted CSP models selected for this study. **Yellow α-helix**, CSP signal peptide 1-26 residues. **Green cartoon**, representative example of lineal-peptide generated by RFdiffusion in complex with full-length CSP. **Red cartoon**, representative example of head-to-tail top cyclic-peptide derived from the **green** lineal peptide in complex with full-length CSP by AFcyclization. In PyMol the cyclic-peptide appeared open when drawn as cartoon, but closed when drawn as sticks.

### ML34.pseML46.pseML53.pse.zip

Representative-top cyclic-peptides coordinates in complex with full-length CSP from three independent-random ML runs (ML-34, ML46, ML53). The *.pse files can be opened in one of the latest versions of PyMol 2024-25.

### Prodigybatch21.py.zip

PyMol/Python script developed to estimate affinities, pdb structures and amino acid contacts of the CSP-peptide complexes. The outputs included the following directories: all_pdb/ (for each of the pdb file complexes), all_contacts/ (a text file containing all predicted amino acid contacts), all_pml (PyMol pml scripts to draw all side chains in the contact surfaces), and all_Kcal (estimated average of total docking affinities in Kcal/mol). The script also predicts a list of all_kcalpM.txt including the calculated picoMolar affinities by the formula 10^12*(exp(Kcal/0.592)), and an ordered list at the orderpdb/ directory (containing all_pdb ordered by the affinities of the generated complexes from all_kcalpM.txt).

### diffusionMain34-6.zip

Colab notebook ipynb filled with last results of RFdiffusion including commonly used contigs: A1-397, pdb: CPS.pdb, hotspot: ““, num_designs:16, add_potential: OK, use_beta_model: OK, num_seqs: 4, mpnn_sampling_temp: 0.3, run_aa: C, use_solubleMPNN: OK, initial_guess: OK, and num_recycles: 6. Includes Driver mount at the first cell and automatically Driver download-save the zipped results at the final cell. Two PyMol/Python scripts (RFana13.py and Prodigybatch21.py) were consecutively run to unzip the results, generate directories and order the lineal-peptides according to their Prodigy affinity estimations, respectively.

### af_cyc_design-JCM10.10batch.zip

Colab notebook ipynb to perform Alphafold2 head-to-tail cyclization (N-C terminal bonds) using the fixbb protocol applied to the lineal-peptides generated from RFdiffusion. Batches of cyclization models (6 per peptide) were generated by modified af_ cyc_design.ipynb ^48^ All the cyclic-peptide *.pdbs were submitted to Prodigy and PyMol as mentioned in the text. The notebook includes the batch design code, Driver mount at the first cell, automatically Driver download-save the zipped directory of results at the final cell, and some of the results from one of the runs.

## Authors’ contributions

JC designed, performed and analyzed the computational work and drafted the manuscript.

## Acknowledgements

Thanks are due to Dr. Bonwin M.J.J. of Bijvoet Center for Biomolecular Research, Utrech, The Netherlands for his initial help with the Prodigy local installation at Windows. Thanks are also due to the participating people at the Slack-Boltz forum (https://boltz-community.slack.com/ssb/redirect) and the Alphafold Discord forum (https://discord.com/channels/). I must thank also to the Gemini /chatGTP maintenance personnel for their help with the developing of multiple Python coding challenges and ipynb modifications.

