## Supplementary material for "*Plasmodium* circumsporozoites targeted by RFdiffusion / cyclic-peptides": af_cyc_designJCM10_10batch.ipynb: - af_cyc_design_JCM10_10batch.ipynb

```
# %% [markdown]
# **#AF_CYC_DESIGN_JCM10.10batch**

**Cyclic peptide structure prediction and design using AlphaFold**

Stephen Rettie, Katelyn Campbell, Asim Bera, Alex Kang, Simon Kozlov, Joshmyn De La Cruz, Victor Adebomi, Guangfeng Zhou, Frank DiMaio, Sergey Ovchinnikov, Gaurav Bhardwaj
doi: https://doi.org/10.1101/2023.02.25.529956

Modified 2025.08.10 by JCM for Driver automatization and batch  


# %% [markdown]
# **1. MOUNT DRIVE**

# %% [code]
from google.colab import drive
drive.mount('/content/drive', force_remount=True) # force remount to ensure connection

# %% [markdown]
# **2. SET UP INSTALL SOFTWARE**

# %% [code]
#@title setup (~2 minutes)
%%time
import os
if not os.path.isdir("params"):
  # get code
  os.system("pip -q install git+https://github.com/sokrypton/ColabDesign.git@v1.1.1")
  # for debugging
  os.system("ln -s /usr/local/lib/python3.*/dist-packages/colabdesign colabdesign")
  # download params
  os.system("mkdir params")
  os.system("apt-get install aria2 -qq")
  os.system("aria2c -q -x 16 https://storage.googleapis.com/alphafold/alphafold_params_2022-12-06.tar")
  os.system("tar -xf alphafold_params_2022-12-06.tar -C params")

import warnings
warnings.simplefilter(action='ignore', category=FutureWarning)
import os
from colabdesign import mk_afdesign_model, clear_mem
from IPython.display import HTML
from google.colab import files
import numpy as np
import jax
import jax.numpy as jnp
from colabdesign.af.alphafold.common import residue_constants

def get_pdb(pdb_code=""):
  if pdb_code is None or pdb_code == "":
    upload_dict = files.upload()
    pdb_string = upload_dict[list(upload_dict.keys())[0]]
    with open("tmp.pdb","wb") as out: out.write(pdb_string)
    return "tmp.pdb"
  elif os.path.isfile(pdb_code):
    return pdb_code
  elif len(pdb_code) == 4:
    os.system(f"wget -qnc https://files.rcsb.org/view/{pdb_code}.pdb")
    return f"{pdb_code}.pdb"
  else:
    os.system(f"wget -qnc https://alphafold.ebi.ac.uk/files/AF-{pdb_code}-F1-model_v3.pdb")
    return f"AF-{pdb_code}-F1-model_v3.pdb"

def add_cyclic_offset(self, offset_type=2):
  #add cyclic offset to connect N and C term
  def cyclic_offset(L):
    i = np.arange(L)
    ij = np.stack([i,i+L],-1)
    offset = i[:,None] - i[None,:]
    c_offset = np.abs(ij[:,None,:,None] - ij[None,:,None,:]).min((2,3))
    if offset_type == 1:
      c_offset = c_offset
    elif offset_type >= 2:
      a = c_offset < np.abs(offset)
      c_offset[a] = -c_offset[a]
    if offset_type == 3:
      idx = np.abs(c_offset) > 2
      c_offset[idx] = (32 * c_offset[idx] )/  abs(c_offset[idx])
    return c_offset * np.sign(offset)
  idx = self._inputs["residue_index"]
  offset = np.array(idx[:,None] - idx[None,:])
  if self.protocol == "binder":
    c_offset = cyclic_offset(self._binder_len)
    offset[self._target_len:,self._target_len:] = c_offset
  if self.protocol in ["fixbb","partial","hallucination"]:
    Ln = 0
    for L in self._lengths:
      offset[Ln:Ln+L,Ln:Ln+L] = cyclic_offset(L)
      Ln += L
  self._inputs["offset"] = offset


#RMSD
def _np_rmsdist(true, pred, use_jax=True):
  _np = jnp if use_jax else np
  d = true[...,None,:] - pred[...,None,:,:]
  d = _np.sqrt(_np.sum(d * d, axis=-1) + 1e-8)
  d = _np.mean(d, axis=(-1, -2)) # mean over (N,N) pairs
  return d
def _np_rmsd(true, pred, use_jax=True):
  _np = jnp if use_jax else np
  p = true - true.mean(-2,keepdims=True)
  q = pred - pred.mean(-2,keepdims=True)
  p = p @ _np_kabsch(p, q, use_jax=use_jax)
  return _np.sqrt(_np.square(p-q).sum(-1).mean(-1) + 1e-8)
def _np_kabsch(a, b, use_jax=True):
  _np = jnp if use_jax else np
  a_centered = a - a.mean(0)
  b_centered = b - b.mean(0)
  # calculate covariance matrix
  C = _np.transpose(a_centered) @ b_centered
  # use SVD to find the rotation matrix
  V, S, W = _np.linalg.svd(C)
  # ensure right-handedness
  d = _np.sign(_np.linalg.det(W @ _np.transpose(V)))
  # calculate rotation matrix
  U = W @ _np.diag([1, 1, d]) @ _np.transpose(V)
  return U
def jnp_rmsdist(true, pred):
  return _np_rmsdist(true, pred)
def jnp_rmsd(true, pred, add_dist=False):
  rmsd = _np_rmsd(true, pred)
  if add_dist: rmsd = (rmsd + _np_rmsdist(true, pred))/2
  return rmsd
def jnp_kabsch_w(a, b, weights):
  return _np_kabsch(a * weights[:,None], b)
def jnp_rmsd_w(true, pred, weights):
  p = true - (true * weights[:,None]).sum(0,keepdims=True)/weights.sum()
  q = pred - (pred * weights[:,None]).sum(0,keepdims=True)/weights.sum()
  p = p @ _np_kabsch(p * weights[:,None], q)
  return jnp.sqrt((weights*jnp.square(p-q).sum(-1)).sum()/weights.sum() + 1e-8)
'''
def add_rmsd_loss(self, weight=20):                           #increase their weight? =>10
  #add rmsd loss
  def loss_fn(inputs, outputs):
    xyz = outputs["structure_module"]
    ca = xyz["final_atom_positions"][:,residue_constants.atom_order["CA"]]
    rmsd = jnp_rmsd(inputs["CA"], ca)
    return {"rmsd":rmsd}
  self._callbacks["model"]["loss"].append(loss_fn)
  self.opt["weights"]["rmsd"] = weight
 '''
def add_rmsd_loss(self, weight=20):                           #increase their weight? =>10
  #add rmsd loss
  def loss_fn(inputs, outputs):
     xyz = outputs["structure_module"]
    # Check if 'CA' key exists in inputs
     if "CA" in inputs:
       ca = xyz["final_atom_positions"][:,residue_constants.atom_order["CA"]]
       rmsd = jnp_rmsd(inputs["CA"], ca)
       return {"rmsd":rmsd}
     else:
      # Handle case where 'CA' key is missing
       print("Warning: 'CA' key not found in inputs. Skipping RMSD loss calculation.")
       return {}  # Return empty dictionary to avoid errors
  self._callbacks["model"]["loss"].append(loss_fn)
  self.opt["weights"]["rmsd"] = weight#rmsd_value=1


# %% [markdown]
# **3. VARIABLES FOR FIXBB PROTOCOL**

# %% [code]
#@title 3. Variables for fixbb protocol
# Define input and output paths
# For batch processing, 'pdb' will now represent the input directory
pdb_input_dir = "/content/drive/MyDrive/ml53pdb" #@param {type:"string"}# Path to the directory containing original PDB files in Google Drive
name = "ml53pdb" #@param {type:"string"} # A base name for the output directory

# Variables for design process
num_models_to_generate = 6 #@param ["1","6", "9", "19", "29"] {type:"raw"}
design_stages = (30, 20, 6) #@param {type:"raw"}
add_cyclic_offset_enabled = True #@param {type:"boolean"}
rmsd_loss_weight = 2 #@param {type:"raw"} #adjust as needed
num_recycles = 3 #@param {type:"raw"}
temperature = 1 #@param {type:"raw"} # Note: Temperature is not directly used by design_3stage, but kept for potential future use or other parts of the code.

# Output directory for the batch results
output_base_dir = f"/content/drive/MyDrive/{name}_output"  # Base directory in Google Drive
os.makedirs(output_base_dir, exist_ok=True)
print (f"OUTPUT_BASE_DIR: ", output_base_dir)

# %% [markdown]
# **4. FIXED BACKBONE DESIGN (FIXBB)**

# %% [code]
#@title 4. Fixed Backbone Design (FIXBB)
import os
from colabdesign import mk_afdesign_model
from colabdesign.shared.protein import  pdb_to_string
import glob # Import the glob module to find files
nn=0
# Get list of PDB files from the input directory
pdb_files = glob.glob(os.path.join(pdb_input_dir, "*.pdb"))

if not pdb_files:
    print(f"No PDB files found in {pdb_input_dir}")
else:
    print(f"Found {len(pdb_files)} PDB files in {pdb_input_dir}. Starting batch processing...")

    for pdb_file in pdb_files:
        # Initialize af_model inside the loop to ensure clean memory state for each PDB
        af_model = mk_afdesign_model(protocol="fixbb")
        nn=nn+1
        # Extract the base name of the PDB file (without extension) to use for output directory
        pdb_name = os.path.splitext(os.path.basename(pdb_file))[0]
        output_dir = os.path.join(output_base_dir, pdb_name)
        os.makedirs(output_dir, exist_ok=True)
        print(f"\n      PROCESSING {pdb_file} number {nn}...")
        print(f"Saving outputs to {output_dir}")


        # Prepare inputs using the current pdb_file
        af_model.prep_inputs(pdb_filename=pdb_file, chain="B")

        # Add cyclic offset and RMSD loss if enabled/specified in the Variables cell
        if add_cyclic_offset_enabled:
            add_cyclic_offset(af_model, offset_type=2)

        add_rmsd_loss(af_model, weight=rmsd_loss_weight) #adjust as needed

        # Run the design process
        for i in range(num_models_to_generate):
            print(f"{nn} - {pdb_name} Generating model {i+1}/{num_models_to_generate}") # ) for {pdb_name}...")
            af_model.design_3stage(*design_stages, num_recycles=num_recycles)
            filepath = os.path.join(output_dir, f"model_{i}.pdb")
            af_model.save_pdb(filepath)

        # Clear memory after processing each PDB file
        clear_mem()
        # Explicitly delete the model and inputs to help release memory
        del af_model
        # You might also want to consider jax.clear_caches() if the issue persists,
        # but re-initializing the model is usually more effective for colabdesign.


    print("\nBatch processing completed.")

# %% [markdown]
# **5. A + B chains**

# %% [code]
#@title 5. A + B chains
import os
from Bio.PDB import PDBParser, PDBIO
from Bio.PDB.Structure import Structure
from Bio.PDB.Model import Model
from Bio.PDB.Chain import Chain
from Bio.PDB.Residue import Residue
from Bio.PDB.Atom import Atom
import glob

# Ensure get_pdb function is available (although not used for original_pdb_input anymore)
# If you ran the setup cell (cell with comment #@title setup (~2 minutes)), this function should be defined.
# If not, you might need to re-run the setup cell.
# from colabdesign.shared.protein import get_pdb # Assuming get_pdb is defined in colabdesign.shared.protein or earlier in the notebook

def extract_chain(pdb_file_path, chain_id):
    """Extracts a specific chain from a PDB file."""
    parser = PDBParser()
    structure = parser.get_structure("structure", pdb_file_path)
    for model in structure:
        for chain in model:
            if chain.get_id() == chain_id:
                return chain
    return None

def combine_pdb_chains(chain_a_structure, chain_b_file, output_file):
    """
    Combines a Bio.PDB Chain object for chain A with a chain from a chain B PDB file
    by creating new Chain objects and copying residues and atoms.
    """
    parser = PDBParser()

    # Load chain B from the generated file
    structure_b = parser.get_structure("structure_b", chain_b_file)
    chain_b_structure = None
    # Assuming the generated PDBs have only one chain
    for model in structure_b:
        for chain in model:
             chain_b_structure = chain # Get the first (and likely only) chain
             break
        if chain_b_structure:
            break

    if not chain_b_structure:
        print(f"Error: Could not extract a chain from {chain_b_file}")
        return

    # Create a new combined structure and model
    combined_structure = Structure("combined")
    target_model = Model(0)
    combined_structure.add(target_model)

    # Create a new Chain object for chain A and copy its residues and atoms
    new_chain_a = Chain("A")
    for residue in chain_a_structure:
        # Create a new residue with the same information
        new_residue_a = Residue(residue.get_id(), residue.get_resname(), residue.get_segid())
        for atom in residue:
            # Create a new atom with the same information
            new_atom_a = Atom(atom.get_name(), atom.get_coord(), atom.get_bfactor(), atom.get_occupancy(), atom.get_altloc(), atom.get_fullname(), atom.get_serial_number(), element=atom.element)
            new_residue_a.add(new_atom_a)
        new_chain_a.add(new_residue_a)
    target_model.add(new_chain_a)

    # Explicitly set chain ID of extracted chain B to a temporary ID
    chain_b_structure.id = 'TEMP'

    # Create a new Chain object with the desired ID 'B'
    new_chain_b = Chain("B")
    for residue in chain_b_structure:
        # Create a new residue with the same information
        new_residue_b = Residue(residue.get_id(), residue.get_resname(), residue.get_segid())
        for atom in residue:
            # Create a new atom with the same information
            new_atom_b = Atom(atom.get_name(), atom.get_coord(), atom.get_bfactor(), atom.get_occupancy(), atom.get_altloc(), atom.get_fullname(), atom.get_serial_number(), element=atom.element)
            # Set a high B-factor for Chain B atoms as a visualization hint
            new_atom_b.set_bfactor(99.99)
            new_residue_b.add(new_atom_b)
        new_chain_b.add(new_residue_b)

    target_model.add(new_chain_b)


    # Save the combined structure
    io = PDBIO()
    io.set_structure(combined_structure)
    io.save(output_file)
    print(f"Combined structure saved to {output_file}")


# Define the input and output directories
# Use the batch input and output directories defined in the Variables cell
# original_pdb_input is no longer a single file, but we need to access original files
# from pdb_input_dir to get Chain A for each batch entry.
generated_pdb_base_dir = output_base_dir # Output from cell 4
combined_pdb_base_dir = f"/content/drive/MyDrive/{name}_combined_pdb" # Base directory to save the combined PDBs

# Create the base output directory for combined PDBs if it doesn't exist
os.makedirs(combined_pdb_base_dir, exist_ok=True)

# Get the list of subdirectories in the generated_pdb_base_dir
# Each subdirectory corresponds to an original PDB file processed in the batch
processed_pdb_dirs = [d for d in os.listdir(generated_pdb_base_dir) if os.path.isdir(os.path.join(generated_pdb_base_dir, d))]

if not processed_pdb_dirs:
    print(f"No processed PDB subdirectories found in {generated_pdb_base_dir}. Please run cell 4 first.")
else:
    print(f"Found {len(processed_pdb_dirs)} processed PDB subdirectories. Starting chain combination...")

    for pdb_name in processed_pdb_dirs:
        original_pdb_file_path = os.path.join(pdb_input_dir, f"{pdb_name}.pdb")
        generated_pdb_dir = os.path.join(generated_pdb_base_dir, pdb_name)
        combined_pdb_dir = os.path.join(combined_pdb_base_dir, pdb_name)

        # Create the output directory for combined PDBs for this specific PDB
        os.makedirs(combined_pdb_dir, exist_ok=True)
        print(f"\nProcessing batch entry for {pdb_name}...")
        print(f"Original PDB: {original_pdb_file_path}")
        print(f"Generated PDBs directory: {generated_pdb_dir}")
        print(f"Saving combined outputs to: {combined_pdb_dir}")


        # Extract chain A from the original PDB file for this batch entry
        print(f"Extracting Chain A from {original_pdb_file_path}...")
        chain_a_structure = extract_chain(original_pdb_file_path, "A")

        if chain_a_structure:
            print("Chain A extracted successfully.")
            # Iterate through the generated chain B files in the subdirectory and combine them with chain A
            print(f"Combining Chain A with generated Chain B models from {generated_pdb_dir}...")
            # Use glob to find the model_*.pdb files in the generated directory
            generated_b_files = glob.glob(os.path.join(generated_pdb_dir, "model_*.pdb"))

            if not generated_b_files:
                print(f"No generated model PDB files found in {generated_pdb_dir}. Skipping combination for {pdb_name}.")
            else:
                for chain_b_file in generated_b_files:
                    filename = os.path.basename(chain_b_file)
                    output_file = os.path.join(combined_pdb_dir, f"combined_{filename}")
                    combine_pdb_chains(chain_a_structure, chain_b_file, output_file)
                print(f"Finished combining PDB files for {pdb_name}.")

        else:
            print(f"Could not extract chain A from {original_pdb_file_path}. Cannot proceed with combining PDBs for {pdb_name}.")

    print("\nBatch chain combination completed.")

# %% [markdown]
# **6. GENERATE BATCHPDB DIR**

# %% [code]
#@title generate batchpdb dir
import os
import shutil
import glob

# Define the base directory where combined PDBs were saved in cell 5
# Ensure this matches the combined_pdb_base_dir defined in cell 5
# We'll reconstruct it based on the 'name' variable from cell 3
# If cell 3's output_base_dir was /content/drive/MyDrive/{name}_output,
# combined_pdb_base_dir was /content/drive/MyDrive/{name}_combined_pdb
#name = "cyclizations_batch" #@param {type:"string"} # Ensure this matches the name used in cell 3
combined_pdb_base_dir = f"/content/drive/MyDrive/{name}_combined_pdb"


# Output folder where all renamed pdb files will be stored in the Colab environment
output_dir = os.path.join("/content/", "batchpdb") # Save to /content/ to keep it local
os.makedirs(output_dir, exist_ok=True)

print(f"Looking for combined PDBs in: {combined_pdb_base_dir}")
print(f"Saving flattened PDBs to: {output_dir}")


# Get the list of subdirectories in the combined_pdb_base_dir
# Each subdirectory corresponds to an original PDB file processed in the batch
processed_pdb_dirs = [d for d in os.listdir(combined_pdb_base_dir) if os.path.isdir(os.path.join(combined_pdb_base_dir, d))]

if not processed_pdb_dirs:
    print(f"No processed PDB subdirectories found in {combined_pdb_base_dir}. Please ensure cell 5 ran successfully.")
else:
    print(f"Found {len(processed_pdb_dirs)} processed PDB subdirectories. Starting flattening...")

    # Loop through each subdirectory (which corresponds to an original PDB name)
    for pdb_name in processed_pdb_dirs:
        subdir_path = os.path.join(combined_pdb_base_dir, pdb_name)

        # Loop through files in the subdir
        # Look for files starting with "combined_model_" and ending with ".pdb"
        combined_files = glob.glob(os.path.join(subdir_path, "combined_model_*.pdb"))

        if not combined_files:
             print(f"No combined_model_*.pdb files found in {subdir_path}. Skipping.")
             continue

        for file_path in combined_files:
            file_name = os.path.basename(file_path)
            # Extract the model number from the filename (e.g., "combined_model_0.pdb" -> "0")
            try:
                parts = file_name.split('_')
                if len(parts) > 2 and parts[-1].endswith(".pdb"):
                     num_part = parts[-1].replace(".pdb", "")
                else:
                     print(f"Warning: Could not parse model number from filename {file_name}. Skipping.")
                     continue
            except Exception as e:
                 print(f"Error parsing filename {file_name}: {e}. Skipping.")
                 continue


            # Construct the new filename: original_pdb_name_model_number.pdb
            new_filename = f"{pdb_name}_{num_part}.pdb"
            dst_path = os.path.join(output_dir, new_filename)

            # Copy the file
            try:
                shutil.copy(file_path, dst_path)
                print(f"Copied {file_path} -> {dst_path}")
            except Exception as e:
                 print(f"Error copying file {file_path} to {dst_path}: {e}")


    print("All files copied to 'batchpdb'")

# %% [markdown]
# **7. SAVE TO DRIVE**

# %% [code]
#@title save batchpdb to drive
import os
import shutil
from google.colab import files

# Define the path to the flattened batchpdb directory created in the previous cell
batchpdb_dir = "/content/batchpdb" # Ensure this matches the output_dir in cell 6

# Define the name for the zip file
zip_filename = f"{name}_batchpdb_files.zip" # Give the zip file a distinct name

# Define a temporary path for the zip file in the Colab environment
temp_zip_path = f"/content/{zip_filename}"

# Change the current directory to the parent of the batchpdb directory
%cd /content/

# Zip the contents of the batchpdb directory
# Use os.path.basename(batchpdb_dir) to zip the directory itself relative to /content/
!zip -r {temp_zip_path} {os.path.basename(batchpdb_dir)}

# Download the zip file from the temporary location
files.download(temp_zip_path)

# Copy the zip file from the temporary location to Google Drive
# Define the destination path in Google Drive
drive_zip_path = f"/content/drive/MyDrive/{zip_filename}"
shutil.copy(temp_zip_path, drive_zip_path)

print(f"SAVED flattened batchpdb files to: {drive_zip_path}")
print(f"Downloaded zip file: {zip_filename}")
```
