## Supplementary material for "*Plasmodium* circumsporozoites targeted by RFdiffusion / cyclic-peptides": diffusionMain34_6.ipynb: - DiffusionMain34_6.ipynb

```
# %% [markdown]
#**RFdiffusionMain34-6**
RFdiffusion is a method for structure generation, with or without conditional information (a motif, target etc). It can perform a whole range of protein design challenges as we have outlined in the RFdiffusion [manuscript](https://www.biorxiv.org/content/10.1101/2022.12.09.519842v2).

**<font color="red">NOTE:</font>** in development,working on adding all the options from the manuscript above.
For **instructions**, see end of Notebook.

See [diffusion_foldcond](https://colab.research.google.com/github/sokrypton/ColabDesign/blob/main/rf/examples/diffusion_foldcond.ipynb) for fold conditioning functionality.See [original version](https://colab.research.google.com/github/sokrypton/ColabDesign/blob/main/rf/examples/diffusion_ori.ipynb) of this notebook (from 31Mar2023).

ed.by JCM 5/VIII/2025 GoogleDrive conection added


# %% [markdown]
# **1. MOUNT DRIVE**

---


# %% [code]
#@title mount drive to colab
from google.colab import drive
drive.mount('/content/drive')
print("Google Drive mounted !")

# %% [markdown]
# **2. SETUP RFDIFFUSION**

---


# %% [code]
#@title setup **RFdiffusion** (~3min)
%%time
import os, time, signal
import sys, random, string, re
if not os.path.isdir("params"):
  os.system("apt-get install aria2")
  os.system("mkdir params")
  # send param download into background
  os.system("(\
  aria2c -q -x 16 https://files.ipd.uw.edu/krypton/schedules.zip; \
  aria2c -q -x 16 http://files.ipd.uw.edu/pub/RFdiffusion/6f5902ac237024bdd0c176cb93063dc4/Base_ckpt.pt; \
  aria2c -q -x 16 http://files.ipd.uw.edu/pub/RFdiffusion/e29311f6f1bf1af907f9ef9f44b8328b/Complex_base_ckpt.pt; \
  aria2c -q -x 16 http://files.ipd.uw.edu/pub/RFdiffusion/f572d396fae9206628714fb2ce00f72e/Complex_beta_ckpt.pt; \
  aria2c -q -x 16 https://storage.googleapis.com/alphafold/alphafold_params_2022-12-06.tar; \
  tar -xf alphafold_params_2022-12-06.tar -C params; \
  touch params/done.txt) &")

if not os.path.isdir("RFdiffusion"):
  print("installing RFdiffusion...")
  os.system("git clone https://github.com/sokrypton/RFdiffusion.git")
  os.system("pip install jedi omegaconf hydra-core icecream pyrsistent pynvml decorator")
  os.system("pip install git+https://github.com/NVIDIA/dllogger#egg=dllogger")
  # 17Mar2024: adding --no-dependencies to avoid installing nvidia-cuda-* dependencies
  os.system("pip install --no-dependencies dgl==2.0.0 -f https://data.dgl.ai/wheels/cu121/repo.html")
  os.system("pip install --no-dependencies e3nn==0.5.5 opt_einsum_fx")
  os.system("cd RFdiffusion/env/SE3Transformer; pip install .")
  os.system("wget -qnc https://files.ipd.uw.edu/krypton/ananas")
  os.system("chmod +x ananas")

if not os.path.isdir("colabdesign"):
  print("installing ColabDesign...")
  os.system("pip -q install git+https://github.com/sokrypton/ColabDesign.git")
  os.system("ln -s /usr/local/lib/python3.*/dist-packages/colabdesign colabdesign")

if not os.path.isdir("RFdiffusion/models"):
  print("downloading RFdiffusion params...")
  os.system("mkdir RFdiffusion/models")
  models = ["Base_ckpt.pt","Complex_base_ckpt.pt","Complex_beta_ckpt.pt"]
  for m in models:
    while os.path.isfile(f"{m}.aria2"):
      time.sleep(5)
  os.system(f"mv {' '.join(models)} RFdiffusion/models")
  os.system("unzip schedules.zip; rm schedules.zip")

if 'RFdiffusion' not in sys.path:
  os.environ["DGLBACKEND"] = "pytorch"
  sys.path.append('RFdiffusion')

from google.colab import files
import json
import numpy as np
import matplotlib.pyplot as plt
from IPython.display import display, HTML
import ipywidgets as widgets
import py3Dmol

from inference.utils import parse_pdb
from colabdesign.rf.utils import get_ca
from colabdesign.rf.utils import fix_contigs, fix_partial_contigs, fix_pdb, sym_it
from colabdesign.shared.protein import pdb_to_string
from colabdesign.shared.plot import plot_pseudo_3D

def get_pdb(pdb_code=None):
  if pdb_code is None or pdb_code == "":
    upload_dict = files.upload()
    pdb_string = upload_dict[list(upload_dict.keys())[0]]
    with open("tmp.pdb","wb") as out: out.write(pdb_string)
    return "tmp.pdb"
  elif os.path.isfile(pdb_code):
    return pdb_code
  elif len(pdb_code) == 4:
    if not os.path.isfile(f"{pdb_code}.pdb1"):
      os.system(f"wget -qnc https://files.rcsb.org/download/{pdb_code}.pdb1.gz")
      os.system(f"gunzip {pdb_code}.pdb1.gz")
    return f"{pdb_code}.pdb1"
  else:
    os.system(f"wget -qnc https://alphafold.ebi.ac.uk/files/AF-{pdb_code}-F1-model_v3.pdb")
    return f"AF-{pdb_code}-F1-model_v3.pdb"

def run_ananas(pdb_str, path, sym=None):
  pdb_filename = f"outputs/{path}/ananas_input.pdb"
  out_filename = f"outputs/{path}/ananas.json"
  with open(pdb_filename,"w") as handle:
    handle.write(pdb_str)

  cmd = f"./ananas {pdb_filename} -u -j {out_filename}"
  if sym is None: os.system(cmd)
  else: os.system(f"{cmd} {sym}")

  # parse results
  try:
    out = json.loads(open(out_filename,"r").read())
    results,AU = out[0], out[-1]["AU"]
    group = AU["group"]
    chains = AU["chain names"]
    rmsd = results["Average_RMSD"]
    print(f"AnAnaS detected {group} symmetry at RMSD:{rmsd:.3}")

    C = np.array(results['transforms'][0]['CENTER'])
    A = [np.array(t["AXIS"]) for t in results['transforms']]

    # apply symmetry and filter to the asymmetric unit
    new_lines = []
    for line in pdb_str.split("\n"):
      if line.startswith("ATOM"):
        chain = line[21:22]
        if chain in chains:
          x = np.array([float(line[i:(i+8)]) for i in [30,38,46]])
          if group[0] == "c":
            x = sym_it(x,C,A[0])
          if group[0] == "d":
            x = sym_it(x,C,A[1],A[0])
          coord_str = "".join(["{:8.3f}".format(a) for a in x])
          new_lines.append(line[:30]+coord_str+line[54:])
      else:
        new_lines.append(line)
    return results, "\n".join(new_lines)

  except:
    return None, pdb_str

def run(command, steps, num_designs=1, visual="none"):

  def run_command_and_get_pid(command):
    pid_file = '/dev/shm/pid'
    os.system(f'nohup {command} & echo $! > {pid_file}')
    with open(pid_file, 'r') as f:
      pid = int(f.read().strip())
    os.remove(pid_file)
    return pid
  def is_process_running(pid):
    try:
      os.kill(pid, 0)
    except OSError:
      return False
    else:
      return True

  run_output = widgets.Output()
  progress = widgets.FloatProgress(min=0, max=1, description='running', bar_style='info')
  display(widgets.VBox([progress, run_output]))

  # clear previous run
  for n in range(steps):
    if os.path.isfile(f"/dev/shm/{n}.pdb"):
      os.remove(f"/dev/shm/{n}.pdb")

  pid = run_command_and_get_pid(command)
  try:
    fail = False
    for _ in range(num_designs):

      # for each step check if output generated
      for n in range(steps):
        wait = True
        while wait and not fail:
          time.sleep(0.1)
          if os.path.isfile(f"/dev/shm/{n}.pdb"):
            pdb_str = open(f"/dev/shm/{n}.pdb").read()
            if pdb_str[-3:] == "TER":
              wait = False
            elif not is_process_running(pid):
              fail = True
          elif not is_process_running(pid):
            fail = True

        if fail:
          progress.bar_style = 'danger'
          progress.description = "failed"
          break

        else:
          progress.value = (n+1) / steps
          if visual != "none":
            with run_output:
              run_output.clear_output(wait=True)
              if visual == "image":
                xyz, bfact = get_ca(f"/dev/shm/{n}.pdb", get_bfact=True)
                fig = plt.figure()
                fig.set_dpi(100);fig.set_figwidth(6);fig.set_figheight(6)
                ax1 = fig.add_subplot(111);ax1.set_xticks([]);ax1.set_yticks([])
                plot_pseudo_3D(xyz, c=bfact, cmin=0.5, cmax=0.9, ax=ax1)
                plt.show()
              if visual == "interactive":
                view = py3Dmol.view(js='https://3dmol.org/build/3Dmol.js')
                view.addModel(pdb_str,'pdb')
                view.setStyle({'cartoon': {'colorscheme': {'prop':'b','gradient': 'roygb','min':0.5,'max':0.9}}})
                view.zoomTo()
                view.show()
        if os.path.exists(f"/dev/shm/{n}.pdb"):
          os.remove(f"/dev/shm/{n}.pdb")
      if fail:
        progress.bar_style = 'danger'
        progress.description = "failed"
        break

    while is_process_running(pid):
      time.sleep(0.1)

  except KeyboardInterrupt:
    os.kill(pid, signal.SIGTERM)
    progress.bar_style = 'danger'
    progress.description = "stopped"

def run_diffusion(contigs, path, pdb=None, iterations=50,
                  symmetry="none", order=1, hotspot=None,
                  chains=None, add_potential=False, partial_T="auto",
                  num_designs=1, use_beta_model=False, visual="none"):

  full_path = f"outputs/{path}"
  os.makedirs(full_path, exist_ok=True)
  opts = [f"inference.output_prefix={full_path}",
          f"inference.num_designs={num_designs}"]

  if chains == "": chains = None

  # determine symmetry type
  if symmetry in ["auto","cyclic","dihedral"]:
    if symmetry == "auto":
      sym, copies = None, 1
    else:
      sym, copies = {"cyclic":(f"c{order}",order),
                     "dihedral":(f"d{order}",order*2)}[symmetry]
  else:
    symmetry = None
    sym, copies = None, 1

  # determine mode
  contigs = contigs.replace(","," ").replace(":"," ").split()
  is_fixed, is_free = False, False
  fixed_chains = []
  for contig in contigs:
    for x in contig.split("/"):
      a = x.split("-")[0]
      if a[0].isalpha():
        is_fixed = True
        if a[0] not in fixed_chains:
          fixed_chains.append(a[0])
      if a.isnumeric():
        is_free = True
  if len(contigs) == 0 or not is_free:
    mode = "partial"
  elif is_fixed:
    mode = "fixed"
  else:
    mode = "free"

  # fix input contigs
  if mode in ["partial","fixed"]:
    pdb_str = pdb_to_string(get_pdb(pdb), chains=chains)
    if symmetry == "auto":
      a, pdb_str = run_ananas(pdb_str, path)
      if a is None:
        print(f'ERROR: no symmetry detected')
        symmetry = None
        sym, copies = None, 1
      else:
        if a["group"][0] == "c":
          symmetry = "cyclic"
          sym, copies = a["group"], int(a["group"][1:])
        elif a["group"][0] == "d":
          symmetry = "dihedral"
          sym, copies = a["group"], 2 * int(a["group"][1:])
        else:
          print(f'ERROR: the detected symmetry ({a["group"]}) not currently supported')
          symmetry = None
          sym, copies = None, 1

    elif mode == "fixed":
      pdb_str = pdb_to_string(pdb_str, chains=fixed_chains)

    pdb_filename = f"{full_path}/input.pdb"
    with open(pdb_filename, "w") as handle:
      handle.write(pdb_str)

    parsed_pdb = parse_pdb(pdb_filename)
    opts.append(f"inference.input_pdb={pdb_filename}")
    if mode in ["partial"]:
      if partial_T == "auto":
        iterations = int(80 * (iterations / 200))
      else:
        iterations = int(partial_T)
      opts.append(f"diffuser.partial_T={iterations}")
      contigs = fix_partial_contigs(contigs, parsed_pdb)
    else:
      opts.append(f"diffuser.T={iterations}")
      contigs = fix_contigs(contigs, parsed_pdb)
  else:
    opts.append(f"diffuser.T={iterations}")
    parsed_pdb = None
    contigs = fix_contigs(contigs, parsed_pdb)

  if hotspot is not None and hotspot != "":
    hotspot = ",".join(hotspot.replace(","," ").split())
    opts.append(f"ppi.hotspot_res='[{hotspot}]'")

  # setup symmetry
  if sym is not None:
    sym_opts = ["--config-name symmetry", f"inference.symmetry={sym}"]
    if add_potential:
      sym_opts += ["'potentials.guiding_potentials=[\"type:olig_contacts,weight_intra:1,weight_inter:0.1\"]'",
                   "potentials.olig_intra_all=True","potentials.olig_inter_all=True",
                   "potentials.guide_scale=2","potentials.guide_decay=quadratic"]
    opts = sym_opts + opts
    contigs = sum([contigs] * copies,[])

  opts.append(f"'contigmap.contigs=[{' '.join(contigs)}]'")
  opts += ["inference.dump_pdb=True","inference.dump_pdb_path='/dev/shm'"]
  if use_beta_model:
    opts += ["inference.ckpt_override_path=./RFdiffusion/models/Complex_beta_ckpt.pt"]

  print("mode:", mode)
  print("output:", full_path)
  print("contigs:", contigs)

  opts_str = " ".join(opts)
  cmd = f"./RFdiffusion/run_inference.py {opts_str}"
  print(cmd)

  # RUN
  run(cmd, iterations, num_designs, visual=visual)

  # fix pdbs
  for n in range(num_designs):
    pdbs = [f"outputs/traj/{path}_{n}_pX0_traj.pdb",
            f"outputs/traj/{path}_{n}_Xt-1_traj.pdb",
            f"{full_path}_{n}.pdb"]
    for pdb in pdbs:
      with open(pdb,"r") as handle: pdb_str = handle.read()
      with open(pdb,"w") as handle: handle.write(fix_pdb(pdb_str, contigs))

  return contigs, copies

# %% [markdown]
# **3. SELECT VARIABLES & RUN TO BACKBONE GENERATION**

---


# %% [code]
%%time
#@title run **RFdiffusion** to generate a backbone
name = "ml53" #@param {type:"string"}
drive_folder_path = "/content/drive/My Drive/ml" #@param {type:"string"}
contigs = "'A1-397:34'" #@param {type:"string"}
pdb = "" #@param {type:"string"}
iterations = 50 #@param ["25", "50", "100", "150", "200"] {type:"raw"}
hotspot = "" #@param {type:"string"}
num_designs = 16 #@param ["1", "2", "4", "8", "16", "32"] {type:"raw"}
visual = "image" #@param ["none", "image", "interactive"]
#@markdown ---
#@markdown **symmetry** settings
#@markdown ---
symmetry = "none" #@param ["none", "auto", "cyclic", "dihedral"]
order = 1 #@param ["1", "2", "3", "4", "5", "6", "7", "8", "9", "10", "11", "12"] {type:"raw"}
chains = "" #@param {type:"string"}
add_potential = True #@param {type:"boolean"}
#@markdown - `symmetry='auto'` enables automatic symmetry dectection with [AnAnaS](https://team.inria.fr/nano-d/software/ananas/).
#@markdown - `chains="A,B"` filter PDB input to these chains (may help auto-symm detector)
#@markdown - `add_potential` to discourage clashes between chains
#@markdown ---
#@markdown **advanced** settings
#@markdown ---
partial_T = "auto" # @param ["auto", "10", "20", "40", "60", "80"]
#@markdown - specify number of noising steps (only used for the partial diffusion protocol)
use_beta_model = True #@param {type:"boolean"}
#@markdown - downnumber helices, better balance.

# determine where to save
path = name
while os.path.exists(f"outputs/{path}_0.pdb"):
  path = name + "_" + ''.join(random.choices(string.ascii_lowercase + string.digits, k=5))

flags = {"contigs":contigs,
         "pdb":pdb,
         "order":order,
         "iterations":iterations,
         "symmetry":symmetry,
         "hotspot":hotspot,
         "path":path,
         "chains":chains,
         "add_potential":add_potential,
         "num_designs":num_designs,
         "use_beta_model":use_beta_model,
         "visual":visual,
         "partial_T":partial_T}

for k,v in flags.items():
  if isinstance(v,str):
    flags[k] = v.replace("'","").replace('"','')

contigs, copies = run_diffusion(**flags)

# %% [markdown]
# **4. DISPLAY 3D STRUCTURES**

---


# %% [code]
#@title Display 3D structure {run: "auto"}
animate = "interactive" #@param ["none", "movie", "interactive"]
color = "chain" #@param ["rainbow", "chain", "plddt"]
denoise = True
dpi = 100 #@param ["100", "200", "400"] {type:"raw"}
from colabdesign.shared.plot import pymol_color_list
from colabdesign.rf.utils import get_ca, get_Ls, make_animation
from string import ascii_uppercase,ascii_lowercase
alphabet_list = list(ascii_uppercase+ascii_lowercase)

def plot_pdb(num=0):
  if denoise:
    pdb_traj = f"outputs/traj/{path}_{num}_pX0_traj.pdb"
  else:
    pdb_traj = f"outputs/traj/{path}_{num}_Xt-1_traj.pdb"
  if animate in ["none","interactive"]:
    hbondCutoff = 4.0
    view = py3Dmol.view(js='https://3dmol.org/build/3Dmol.js')
    if animate == "interactive":
      pdb_str = open(pdb_traj,'r').read()
      view.addModelsAsFrames(pdb_str,'pdb',{'hbondCutoff':hbondCutoff})
    else:
      pdb = f"outputs/{path}_{num}.pdb"
      pdb_str = open(pdb,'r').read()
      view.addModel(pdb_str,'pdb',{'hbondCutoff':hbondCutoff})
    if color == "rainbow":
      view.setStyle({'cartoon': {'color':'spectrum'}})
    elif color == "chain":
      for n,chain,c in zip(range(len(contigs)),
                              alphabet_list,
                              pymol_color_list):
          view.setStyle({'chain':chain},{'cartoon': {'color':c}})
    else:
      view.setStyle({'cartoon': {'colorscheme': {'prop':'b','gradient': 'roygb','min':0.5,'max':0.9}}})
    view.zoomTo()
    if animate == "interactive":
      view.animate({'loop': 'backAndForth'})
    view.show()
  else:
    Ls = get_Ls(contigs)
    xyz, bfact = get_ca(pdb_traj, get_bfact=True)
    xyz = xyz.reshape((-1,sum(Ls),3))[::-1]
    bfact = bfact.reshape((-1,sum(Ls)))[::-1]
    if color == "chain":
      display(HTML(make_animation(xyz, Ls=Ls, dpi=dpi, ref=-1)))
    elif color == "rainbow":
      display(HTML(make_animation(xyz, dpi=dpi, ref=-1)))
    else:
      display(HTML(make_animation(xyz, plddt=bfact*100, dpi=dpi, ref=-1)))


if num_designs > 1:
  output = widgets.Output()
  def on_change(change):
    if change['name'] == 'value':
      with output:
        output.clear_output(wait=True)
        plot_pdb(change['new'])
  dropdown = widgets.Dropdown(
      options=[(f'{k}',k) for k in range(num_designs)],
      value=0, description='design:',
  )
  dropdown.observe(on_change)
  display(widgets.VBox([dropdown, output]))
  with output:
    plot_pdb(dropdown.value)
else:
  plot_pdb()

# %% [markdown]
# **5. RUN MPNN+ALPHAFOLD TO GENERATE SEQUENCE DESIGNS**

---


# %% [code]
%%time
#@title run **ProteinMPNN** to generate a sequence and **AlphaFold** to validate
#@markdown ProteinMPNN Settings
num_seqs = 4 #@param ["1", "2", "4", "8", "16", "32", "64"] {type:"raw"}
mpnn_sampling_temp = 0.3 #@param ["0.0001", "0.1", "0.15", "0.2", "0.25", "0.3", "0.5", "1.0"] {type:"raw"}
rm_aa = "C" #@param {type:"string"}
use_solubleMPNN = True #@param {type:"boolean"}
#@markdown - `mpnn_sampling_temp` - control diversity of sampled sequences. (higher = more diverse).
#@markdown - `rm_aa='C'` - do not use [C]ysteines.
#@markdown - `use_solubleMPNN` - use weights trained only on soluble proteins. See [preprint](https://www.biorxiv.org/content/10.1101/2023.05.09.540044v2).
#@markdown
#@markdown AlphaFold Settings
initial_guess = True #@param {type:"boolean"}
#@markdown - soft initialization with desired coordinates, see [paper](https://www.nature.com/articles/s41467-023-38328-5).
num_recycles = 6 #@param ["0", "1", "2", "3", "6", "12"] {type:"raw"}
#@markdown - for **binder** design, we recommend `initial_guess=True num_recycles=3`
use_multimer = False #@param {type:"boolean"}
#@markdown - `use_multimer` - use AlphaFold Multimer v3 params for prediction.

if not os.path.isfile("params/done.txt"):
  print("downloading AlphaFold params...")
  while not os.path.isfile("params/done.txt"):
    time.sleep(5)

contigs_str = ":".join(contigs)
opts = [f"--pdb=outputs/{path}_0.pdb",
        f"--loc=outputs/{path}",
        f"--contig={contigs_str}",
        f"--copies={copies}",
        f"--num_seqs={num_seqs}",
        f"--num_recycles={num_recycles}",
        f"--rm_aa={rm_aa}",
        f"--mpnn_sampling_temp={mpnn_sampling_temp}",
        f"--num_designs={num_designs}"]
if initial_guess: opts.append("--initial_guess")
if use_multimer: opts.append("--use_multimer")
if use_solubleMPNN: opts.append("--use_soluble")
opts = ' '.join(opts)
!python colabdesign/rf/designability_test.py {opts}

# %% [markdown]
# **6. DISPLAY ~BEST SEQUENCE RESULT**

---


# %% [code]
#@title Display best sequence result
import py3Dmol
def plot_pdb(num = "best"):
  if num == "best":
    with open(f"outputs/{path}/best.pdb","r") as f:
      # REMARK 001 design {m} N {n} RMSD {rmsd}
      info = f.readline().strip('\n').split()
    num = info[3]
  hbondCutoff = 4.0
  view = py3Dmol.view(js='https://3dmol.org/build/3Dmol.js')
  pdb_str = open(f"outputs/{path}_{num}.pdb",'r').read()
  view.addModel(pdb_str,'pdb',{'hbondCutoff':hbondCutoff})
  pdb_str = open(f"outputs/{path}/best_design{num}.pdb",'r').read()
  view.addModel(pdb_str,'pdb',{'hbondCutoff':hbondCutoff})

  view.setStyle({"model":0},{'cartoon':{}}) #: {'colorscheme': {'prop':'b','gradient': 'roygb','min':0,'max':100}}})
  view.setStyle({"model":1},{'cartoon':{'colorscheme': {'prop':'b','gradient': 'roygb','min':0,'max':100}}})
  view.zoomTo()
  view.show()

if num_designs > 1:
  def on_change(change):
    if change['name'] == 'value':
      with output:
        output.clear_output(wait=True)
        plot_pdb(change['new'])
  dropdown = widgets.Dropdown(
    options=["best"] + [str(k) for k in range(num_designs)],
    value="best",
    description='design:',
  )
  dropdown.observe(on_change)
  output = widgets.Output()
  display(widgets.VBox([dropdown, output]))
  with output:
    plot_pdb(dropdown.value)
else:
  plot_pdb()

# %% [markdown]
# **7. ZIP AND DOWNLOAD**

---


# %% [code]
#@title Package and download results
!zip -r {path}.result.zip outputs/{path}* outputs/traj/{path}*
files.download(f"{path}.result.zip")
# Export to Google Drive
import os
import shutil
from datetime import datetime
from google.colab import files
from pathlib import Path
# Define the source file path
source_file_path = f"/content/{path}.result.zip" # Update this path if needed
#drive_folder_path = "/content/drive/My Drive/Lm30" #@param {type:"string"}
target_filename = f"{path}.result.zip"
# Ensure the target directory exists in Google Drive
target_dir = Path(drive_folder_path)
target_dir.mkdir(parents=True, exist_ok=True)
print(f"Saving file to: {target_dir / target_filename}")
try:
    shutil.copy(source_file_path, target_dir / target_filename)
    print(f"Successfully copied {source_file_path} to {target_dir / target_filename}")
except FileNotFoundError:
    print(f"Error: Source file not found at {source_file_path}")
except Exception as e:
    print(f"Error copying file to Google Drive: {e}")

# %% [markdown]
**Instructions**
Use `contigs` to define continious chains. Use a `:` to define multiple contigs and a `/` to define mutliple segments within a contig.For example:
**unconditional**
- `contigs='100'` - diffuse **monomer** of length 100
- `contigs='50:100'` - diffuse **hetero-oligomer** of lengths 50 and 100
- `contigs='50'` `symmetry='cyclic'` `order=2` - make two copies of the defined contig(s) and add a symmetry constraint, for **homo-oligomeric** diffusion.
**binder design**
- `contigs='A:50'` `pdb='4N5T'` - diffuse a **binder** of length 50 to chain A of defined PDB.
- `contigs='E6-155:70-100'` `pdb='5KQV'` `hotspot='E64,E88,E96'` - diffuse a **binder** of length 70 to 100 (sampled randomly) to chain E and defined hotspot(s).
**motif scaffolding**
 - `contigs='40/A163-181/40'` `pdb='5TPN'`
 - `contigs='A3-30/36/A33-68'` `pdb='6MRR'` - diffuse a loop of length 36 between two segments of defined PDB ranges.
**partial diffusion**
- `contigs=''` `pdb='6MRR'` - noise all coordinates
- `contigs='A1-10'` `pdb='6MRR'` - keep first 10 positions fixed, noise the rest
- `contigs='A'` `pdb='1SSC'` - fix chain A, noise the rest
*hints and tips*
- `pdb=''` leave blank to get an upload prompt
- `contigs='50-100'` use dash to specify a range of lengths to sample from
```
