## Supplementary figures and images for "*Plasmodium* circumsporozoites targeted by RFdiffusion / cyclic-peptides"

### GraphycalAbstract.png

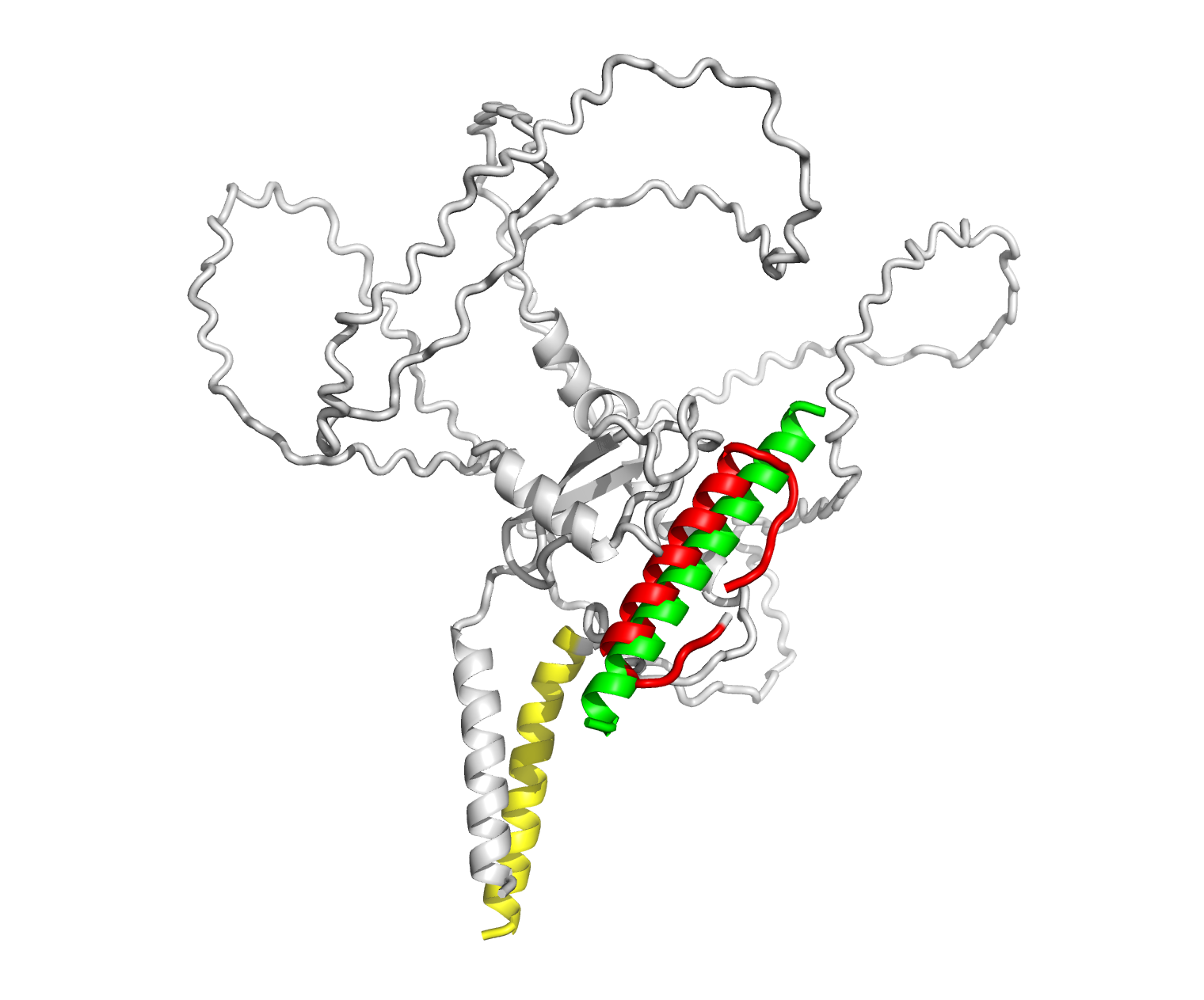
