## Supplementary material for "*Plasmodium* circumsporozoites targeted by RFdiffusion / cyclic-peptides": Prodigybatch21: prodigybatch21.py

```
import os
import os.path
import subprocess
import shutil
import zipfile
import glob
import random
import shutil
import math
from pymol import cmd

#1.PATHS & DELETING "DESIGN" OF PDB-ALL FILES AT CURRENT DIR ############################################################################
current_dir = os.getcwd()
print("Current dir:", current_dir)
prodigy_exe = r"C:\Users\JCOLL\anaconda3\envs\PRODIGY\Scripts\prodigy.exe"
#output_summary = "kcal.txt"
for filename in os.listdir(current_dir):
    if filename.startswith("design"):
        os.rename(filename, filename[6:])

#2. RUN PRODIGY.EXE #############################################################################################################################
for filename in os.listdir(current_dir):
    if filename.endswith(".pdb"):
        print(f"prodigying {filename} ...")
        abs_pdb = os.path.join(current_dir, filename)
        result_file = filename.replace(".pdb", "- prodigy.txt")
        with open(result_file, "w", encoding="utf-8") as outfile:
            startupinfo = subprocess.STARTUPINFO()
            startupinfo.dwFlags |= subprocess.STARTF_USESHOWWINDOW
            subprocess.run([prodigy_exe, abs_pdb, "--selection", "A", "B", "--distance-cutoff", "5.5","--contact_list","--pymol_selection","-q"],
                    startupinfo=startupinfo,stdout=outfile,stderr=subprocess.STDOUT)

#3 TXTS TO ALL_dir:: all_Kcal, all_pdbs, all_pdbs2, all_pml, all_contacts #################################################################################################################################################################################################################
#prodigy kcal dir -q--------------------------------------
nn=0
dst = os.path.join(current_dir, "all_Kcal")
if not os.path.isdir(dst):
     os.mkdir(dst)
for filename in os.listdir(current_dir):
    if filename.endswith("prodigy.txt"):
        nn=nn+1
        src = os.path.join(current_dir,filename)
        dst = os.path.join(current_dir, "all_Kcal")
        shutil.copy(src, dst)
print(str(nn) + " Kcals -->  all_Kcal")

#all_pdbs dir-------------------------------------------
nn=0
dst = os.path.join(current_dir, "all_pdb")
if not os.path.isdir(dst):
     os.mkdir(dst)
for filename in os.listdir(current_dir):
    if filename.endswith(".pdb"):
        nn=nn+1
        src = os.path.join(current_dir,filename)
        dst = os.path.join(current_dir, "all_pdb")
        shutil.copy(src, dst)
print(str(nn) + " pdbs -->  all_pdb")

#all_pml dir------------------------------------------
nn=0
dst = os.path.join(current_dir, "all_pml")
if not os.path.isdir(dst):
     os.mkdir(dst)
for filename in os.listdir(current_dir):
    if filename.endswith(".pml"):
        nn=nn+1
        src = os.path.join(current_dir,filename)
        dst = os.path.join(current_dir, "all_pml")
        shutil.copy(src, dst)
print(str(nn) + " pml -->  all_pml")

#all_contacts dir------------------------------------
dst = os.path.join(current_dir, "all_contacts")
if not os.path.isdir(dst):
     os.mkdir(dst)
for filename in os.listdir(current_dir):
    if filename.endswith(".ic"):
        nn=nn+1
        src = os.path.join(current_dir,filename)
        dst = os.path.join(current_dir, "all_contacts")
        shutil.copy(src, dst)
print(str(nn) + " ic -->  all_contacts")

#deleting extra filenames-------------------------------------
import glob, os, os.path
filelist = glob.glob(os.path.join(current_dir, "*.*"))
filtered_files = [file for file in filelist if not file.endswith(".py")]
for filename in filtered_files:
    os.remove(filename)
print("prodigy predicted all_Kcal, all_pml, all_contacts, all_pdb, all_pdb2 in one dir each")

#4 ALL_TXT TO ALL_KCLPM#####################################################################
kcal_dir = "all_Kcal"
output_file = "all_kcalpM.txt"
data = []
for fname in os.listdir(kcal_dir):
    if fname.endswith(".txt"):
        path = os.path.join(kcal_dir, fname)
        with open(path, "r", encoding="utf-8", errors="ignore") as f:
            lines = f.readlines()
            if len(lines) >= 2:
                second_line = lines[1].strip()
                parts = second_line.split()
                if len(parts) == 2:
                    label, kcal_str = parts
                    try:
                        kcal = float(kcal_str)
                        pM = 1e12 * math.exp(kcal / 0.592)
                        data.append((label, kcal, pM))
                    except ValueError:
                        pass  # skip if kcal_str is not a float
data.sort(key=lambda x: x[2], reverse=False)
with open(output_file, "w") as out:
    for label, kcal, pM in data:
        out.write(f"{label}\t{kcal:.2f}\t{pM:.3f}\n")
print(f"Affinity data written to: {output_file}")

#5. ALL_KCLPM + ALL_PDB TO BATCHPDB #########################################################################################################################
import os
import shutil
current_dir = os.getcwd()
batchpdb_dir = current_dir #os.path.dirname(os.path.abspath(__file__)) # Get the directory where the script is located
order_file_path = os.path.join(batchpdb_dir, "all_kcalpM.txt")
pdb_dir_path = os.path.join(batchpdb_dir, "all_pdb") # The directory containing the original PDB files
ordered_pdb_dir = os.path.join(batchpdb_dir, "ordered_pdb") #batchpd_dir") #ordered_pdb") # New directory for the ordered PDBs
os.makedirs(ordered_pdb_dir, exist_ok=True)
print(f"order from: {order_file_path}") # ordered all_kcalpM.txt 
print(f"PDBs in: {pdb_dir_path}") #get the all_pdb dir
print(f"ordered PDBs to: {ordered_pdb_dir}")
ordered_filenames = []
try:
    with open(order_file_path, 'r') as f: #open read all_kcalpM.txt
        for line in f:
            parts = line.split() #first part of the line
            if parts:
                filename = parts[0] + ".pdb" # Extract the base filename (e.g., "6_n3_0") and pdb
                ordered_filenames.append(filename)
except FileNotFoundError:
    print(f"Error: Order file not found at {order_file_path}")
    exit() # Exit the script if the order file is not found
except Exception as e:
    print(f"Error reading order file: {e}")
    exit()
if not ordered_filenames:
    print("No filenames found in the order file. Exiting.")
else:
    print(f"Found {len(ordered_filenames)} filenames in the order file.")
    print("")
    print("Reordering and copying PDB files...")
    for i, filename in enumerate(ordered_filenames):
        original_pdb_path = os.path.join(pdb_dir_path, filename)
        new_filename = f"{i+1:03d}_{filename}" # Create the new filename with a numerical prefix (e.g., 001_filename.pdb)
        destination_pdb_path = os.path.join(ordered_pdb_dir, new_filename)
        # Check if the original PDB file exists before copying
        if os.path.exists(original_pdb_path):
            try:
                shutil.copy(original_pdb_path, destination_pdb_path)
                # print(f"Copied {original_pdb_path} to {destination_pdb_path}")
            except Exception as e:
                print(f"Error copying file {original_pdb_path} to {destination_pdb_path}: {e}")
        else:
            print(f"Warning: Original PDB file not found: {original_pdb_path}. Skipping.")
    print(f"\nFINISHED PDB REORDERING AT: {ordered_pdb_dir}")
```
